# Deep-learning-enabled morphodynamic analysis of drug responses in a biomimetic fibrin-based 3D glioblastoma invasion model

**DOI:** 10.64898/2026.03.24.713307

**Authors:** Zhipeng Dong, Satvik Kethireddy, Daehyung Kim, Patrick Ting, Batchu Lal, Kwonmoo Lee, Deok-Ho Kim, Eun Hyun Ahn

## Abstract

Glioblastoma (GBM) lethality arises from aggressive invasion and diffuse infiltration of brain tissue. Conventional GBM preclinical models often fail to predict clinical therapeutic efficacy because they do not recapitulate the pathological extracellular matrix (ECM) cues that drive tumor invasion. Here, we present an ECM mimetic 3D platform using a fibrin scaffold to recapitulate the hemorrhagic, pro-thrombotic tumor microenvironment characteristic of high-grade gliomas. This fibrin scaffold induces a pro-invasive phenotype in GBM spheroids by upregulating proliferation/cell cycle- (*MYC, FOXOM1, CCND1*) and invasion-associated-(*CTSS, FOXM1, CCND1*) genes. Traditional cell morphology quantification methods (e.g., circularity) distil complex shapes into singular metrics and cannot capture the nuances of invasion. To address this limitation, we have applied a deep-learning segmentation pipeline (MARS-Net) and high-content morphodynamic descriptors. By using the Preserving Heterogeneity (PHet) algorithm, the 3D platform accurately classifies invasiveness levels and captures the invasion-inhibitory effects of potential repurposable drug candidates. We demonstrate that our model can predict a spheroid’s long-term invasive fate with high accuracy using only partial image sets from early time-points, rather than the complete time-course images. Our work presents an *in vivo*-like, scalable 3D platform integrated with a quantitative high-throughput pipeline to elucidate GBM invasion mechanisms and to evaluate anti-invasive compounds.

## 1. Introduction

Glioblastoma multiforme (GBM) remains one of the most lethal and aggressive type of brain tumors, with a median patient survival time of only 15 months, with less than 5% of patients surviving 5 years.^[1]^ GBM aggressively invades into the local brain tissues and along the vasculature, making it difficult to completely resect the tumor and leading to near-universal recurrence.^[2,3]^ GBM tumors are highly heterogenous and resistant to standard-of-care treatments, including radiation and the alkylating chemotherapeutic agent temozolomide (TMZ).^[4]^ The blood-brain barrier (BBB) further complicates treatment by limiting the penetration of many potential therapeutic agents.^[5]^ Currently, TMZ remains the only FDA-approved first-line chemotherapy for GBM and is the primary FDA-approved chemotherapeutic drug for GBM. Although it exhibits improved central nervous system (CNS) penetration compared to other agents, its delivery is restricted by the BBB efflux transporters and suboptimal polar surface area. Additionally, clinical efficacy is severely compromised by resistance occurring in approximately 50% of patients. The TMZ treatment is effective in tumors with a methylated and silenced MGMT promoter, however it often fails to eliminate the tumor cells, leading to recurrence at the original site in most cases (70-90%).^[6]^

Current preclinical screenings for GBM therapies face significant hurdles, marked by high failure rates of drug candidates in clinical trials. This can be attributed to initial testing in conventional *in vitro* models, such as 2D cell cultures, which lack a tumor microenvironment, Such testing fails to accurately predict how drugs will perform in patients.^[7]^ Human-sourced, 3D tumor spheroid/organoid models can address this critical gap. Unlike 2D cell cultures, as GBM spheroids/organoids can better recapitulate the heterogeneity of cell populations within GBM, while reproducing the 3D microenvironment and cell-cell interaction.^[8]^ By culturing GBM spheroids within biomaterials engineered to mimic the specific pathological extracellular matrix, these models provide a unique opportunity for the direct observation and quantification of GBM cell invasion, a key driver of the disease’s lethality. Furthermore, 3D spheroid/organoid model offers a platform with intermediate complexity, potentially more amenable to higher-throughput screening than complex animal models, thus improving the predictive power of early-stage drug evaluation.^[9]^

In preclinical models, it is important to recapitulate the specific and relevant tumor microenvironment (TME), since the extracellular matrix (ECM) around the tumor plays an important role in mediating tumor invasion and survival.^[10,11]^ The normal brain ECM largely composed of hyaluronan, chondroitin sulfate proteoglycans, and tenascin-R, which intrinsically resists the infiltration of cells.^[12,13]^ However, as gliomas advance, the tumor cells undergo transcriptional activation that leads to the deposition of fibrillar ECM proteins, including fibronectin, collagen, and fibrin.^[14]^ This transformation of the brain’s supportive scaffolding into an adhesive, fibrillar matrix is strongly correlated with decreased patient survival, highlighting the necessity of accurately modeling this altered ECM in biofabricated systems.^[12]^

Fibrin has gained increasing attention as an important biochemical cue in the pro-invasion GBM niche. Excessive angiogenesis within GBM is often associated with hemorrhage and thrombosis, creating an ECM microenvironment rich in fibrin.^[15]^ Intravascular thrombosis, representing local clot formation within tumor vessels, was observed in 45% of primary GBM resections.^[16]^ Fibrin deposition is elevated in GBM tissue compared to fibrin-free normal brain tissue.^[14,15]^ In 3D culture, upregulation of the stem cell markers and promoted proliferation are observed when GBM is cultured in fibrin, effects that are not observed when cultured in traditional 3D matrices such as Matrigel.^[15]^ These findings suggest that fibrin deposition provides an ECM niche that promotes the proliferation and invasion of GBM, especially around the brain vasculature. Consequently, fibrinogen-based hydrogel serves not merely as a passive scaffold, but as an engineered biomaterial capable of instructing the invasive phenotype. In this study, we modeled the local invasion of GBM spheroids within a 3D fibrin scaffold and investigated the mechanism of this specific tumor-ECM interaction.

The high-content functional readout from the preclinical model is key to the predictive power of the system. Most existing 3D tumor spheroid/organoid assays focus on evaluating the viability and proliferation of tumor cells in an ECM-free environment.^[17]^ While some methods address invasive analysis of tumor spheroids, they usually require specialized devices or rely on the measurement of simple morphological features, such as changes in area or diameter.^[18,19]^ Such simple quantitation of tumor invasiveness limits the sensitivity of the assay, highlighting the need for measuring more informative quantitative metrics. The morphodynamic features comprehensively characterize all aspects of the spheroid’s shape, size and motility during invasion. In this study, we have developed an end-to-end, integrated platform for GBM invasion analysis. We have established a 3D GBM invasion model embedded in a biomimetic fibrin matrix and coupled it with a DL-based segmentation and high-content analysis method. Our analytical pipeline utilizes MARS-Net, a deep-learning based segmentation algorithm, for efficient and accurate tumor spheroid morphology tracking from live cell microscopy.^[20]^ The subsequent feature extraction and prioritization algorithm, known as Preserving Heterogeneity (PHet), enables in-depth analysis of the morphodynamic features of invasive tumors spheroids, leading to sensitive detection of different invasive patterns.^[21]^ This synergistic evolution between model complexity and analytical power is crucial for fully realizing the potential of 3D invasion assays in both fundamental research and high-throughput drug discovery.

Using this integrated platform, we evaluated the therapeutic potential capability of drugs, identified from a large-scale drug repurposing screening study.^[22]^ Drug repurposing, the process of finding new uses for existing medications, offers significant strategic advantages over traditional *de novo* drug discovery. Because these drugs have already passed prior safety and toxicity testing, researchers can bypass early-stage clinical trials.^[23]^ This strategy substantially reduces research costs and development timeframes, facilitating faster clinical translation for lethal conditions like GBM where time is a critical factor for patient outcomes. We have demonstrated that four selected drugs demonstrated greater anti-invasion and anti-growth efficacy than the FDA-approved first-line GBM chemotherapy agent temozolomide (TMZ) in our GBM spheroids embedded in3D fibrin scaffolds. Our work presents an *in vivo*-like pre-clinical platform that could facilitate the discovery of anti-invasive and anti-growth therapeutic agents for GBM.

We have hypothesized that a fibrin-rich extracellular matrix (ECM) would better recapitulate the pro-invasive tumor microenvironment compared to Matrigel, a basement membrane extract commonly used for 3D cell culture. To validate this hypothesis, we pursued the following aims: a) establish a 3D invasion platform suitable for spheroids/organoids generated from GBM PDX cells embedded in a fibrin-based hydrogel; b) identify target genes and transcriptomic signatures associated with the fibrin-driven invasive phenotype; and c) quantitatively profile GBM spheroid invasiveness using a deep-learning-based segmentation coupled with high-content analysis **(Figure 1A)**.

**Figure 1.**
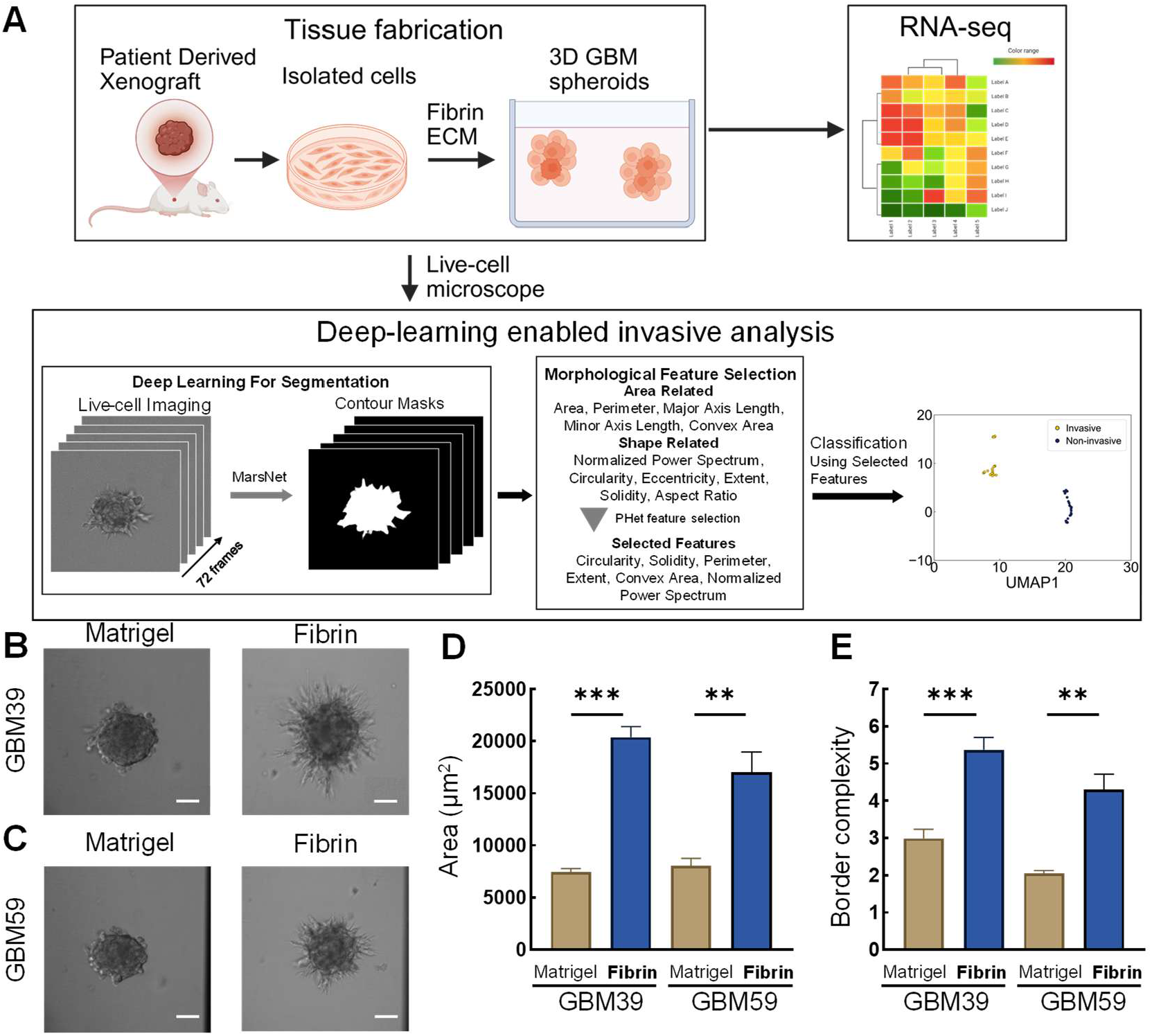
Characterization of GBM invasiveness in 3D fibrin scaffolds. **(A)** Schematic illustration of the study. Representative images of **(B)** GBM PDX 39 and **(C)** GBM PDX 59 spheroids embedded in Matrigel and fibrin scaffolds at 12 hours post-seeding. Scale bars: 200 µm. Quantitative comparison of **(D)** total spheroid area and **(E)** border complexity, defined as the inverse of circularity, of GBM PDX 39 and 59 spheroids cultured in fibrin and Matrigel scaffolds. Statistically significant differences are indicated (\*\**p* < 0.01, \*\*\**p* < 0.001, by the Mann-Whitney U test).

## 2. Results

### 2.1. Biomimetic fibrin scaffold promotes a highly invasive phenotype in GBM spheroids

Live-cell imaging was conducted to investigate morphological differences in patient derived xenograft (PDX) GBM39 and GBM59 spheroids embedded in two distinct 3D hydrogels, fibrin and Matrigel in a high-throughout 96-well format. Spheroids cultured in Matrigel remained a compact, spherical morphology with minimal cell dissemination over the 12-hour culture period. In contrast, spheroids embedded in the fibrin matrix displayed rapid and aggressive invasion, characterized by extensive, irregular protrusions of cells into the surrounding hydrogel **(Figure 1B and C)**. These morphological changes were quantified with total spheroid area and border complexity. Total spheroid areas were significantly higher in GBM spheroids cultured in the fibrin scaffold compared to those in Matrigel **(Figure 1D)**. Furthermore, we quantified the spheroid boundary complexity using the inverse of circularity as an indicator of invasive activity; spheroids in fibrin manifested significantly more complex and irregular borders than their Matrigel-cultured counterparts **(Figure 1E)**. These results indicate that a fibrin-based ECM matrix provides a pro-invasive microenvironment, validating its use as a more relevant model for studying GBM invasion than Matrigel-matrix.

### 2.2. Fibrin induces a pro-proliferative and pro-invasive transcriptomic signature in GBM spheroids

Having established that fibrin promotes an invasive phenotype, we next sought to uncover the underlying molecular mechanisms. We performed bulk RNA sequencing (RNA-seq) on GBM39 spheroids cultured in fibrin for 2 days and compared their transcriptomic profiles to pre-invasive spheroids on day 0 and those cultured in Matrigel on day 2. Principal Component Analysis (PCA) revealed distinct clustering of the samples, indicating that the fibrin scaffold induces a unique transcriptomic state **(Figure 2A)**. The distance matrix of PCA and hierarchical clustering based on the expression further confirms that samples from each condition cluster together **(Figure 2B and C)**.

**Figure 2.**
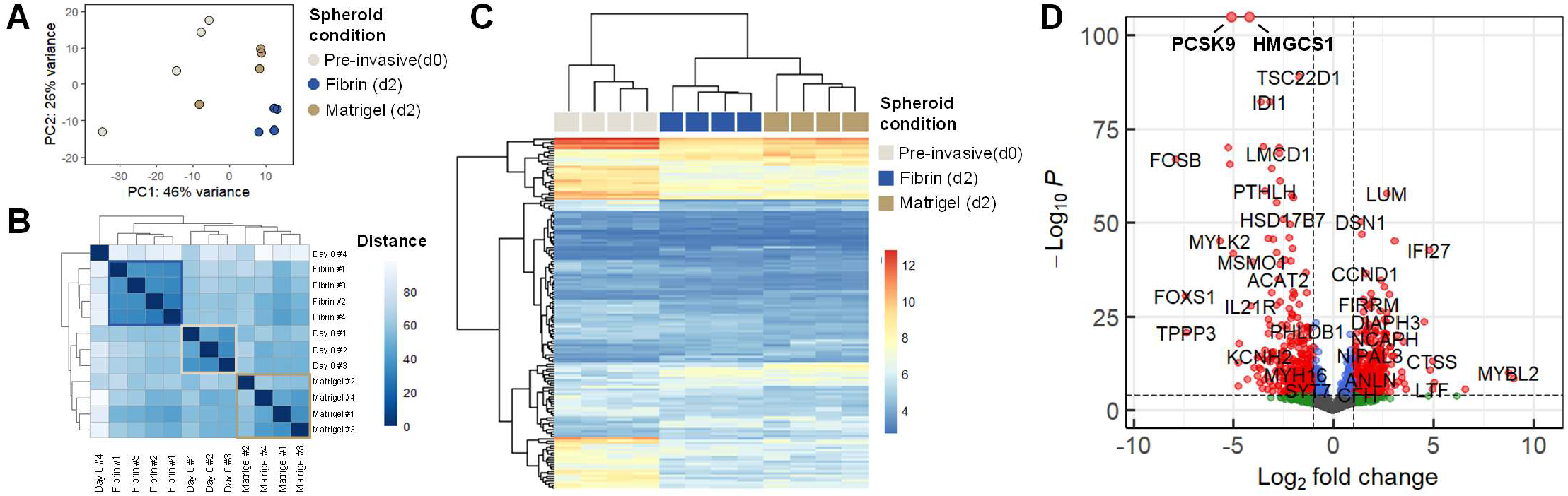
Bulk RNAseq analysis of GBM spheroids in fibrin and Matrigel scaffolds. **(A)** Principal component analysis (PCA) of GBM PDX 39 spheroids across three conditions: day 0 (pre-invasive), day 2 in Matrigel, and day 2 in fibrin. **(B)** Sample distance matrix representing global transcriptomic similarities between samples from the same condition within the PCA analysis. **(C)** Hierarchical clustering of DEGs exhibiting at least two-fold change in expression. **(D)** Volcano plot illustrates DEGs between day 2 fibrin-embedded spheroids and day 0 pre-invasive controls.

We identified a set of differentially expressed genes (DEGs) that were significantly up- and down-regulated in the invasive, fibrin-cultured spheroids **(Figure 2D)**. Network analysis of these DEGs highlighted the significant enrichment of pathways, functions, and regulators related to cell motility, and tumor invasion **(Figure 3A, B)**. Specifically, within these motility-associated networks, we identified a pronounced activation of formin-mediated actin assembly via RHO GTPase signaling cascades, thereby promoting the cytoskeletal remodeling required to drive active cell migration. Notably, we also observed increased expression of *CTSS*, a gene encoding the ECM-remodeling enzyme Cathepsin S, which is involved in the process of tumor migration and invasion **(Figure 3C)**. Concurrently, major pathways involved in cell cycle progression, proliferation, and DNA damage response were activated, while apoptosis pathways were inhibited. Key regulators of the cell cycle, including *FOXM1, MYC*, and *CCND1*, were significantly upregulated in the fibrin environment and considered to drive the observed phenotype **(Figure 3D-F)**. Based on TCGA survival data, patients with high *CTSS, FOXM1*, and *MYC* expression have poor survival outlook, indicating that these markers correlate with advanced stage of GBM and higher lethality. **(Figure 3G-I)** These findings demonstrate that the fibrin matrix actively rewires the molecular profile of GBM cells, promoting a pro-proliferative and pro-invasive transcriptomic signature.

**Figure 3.**
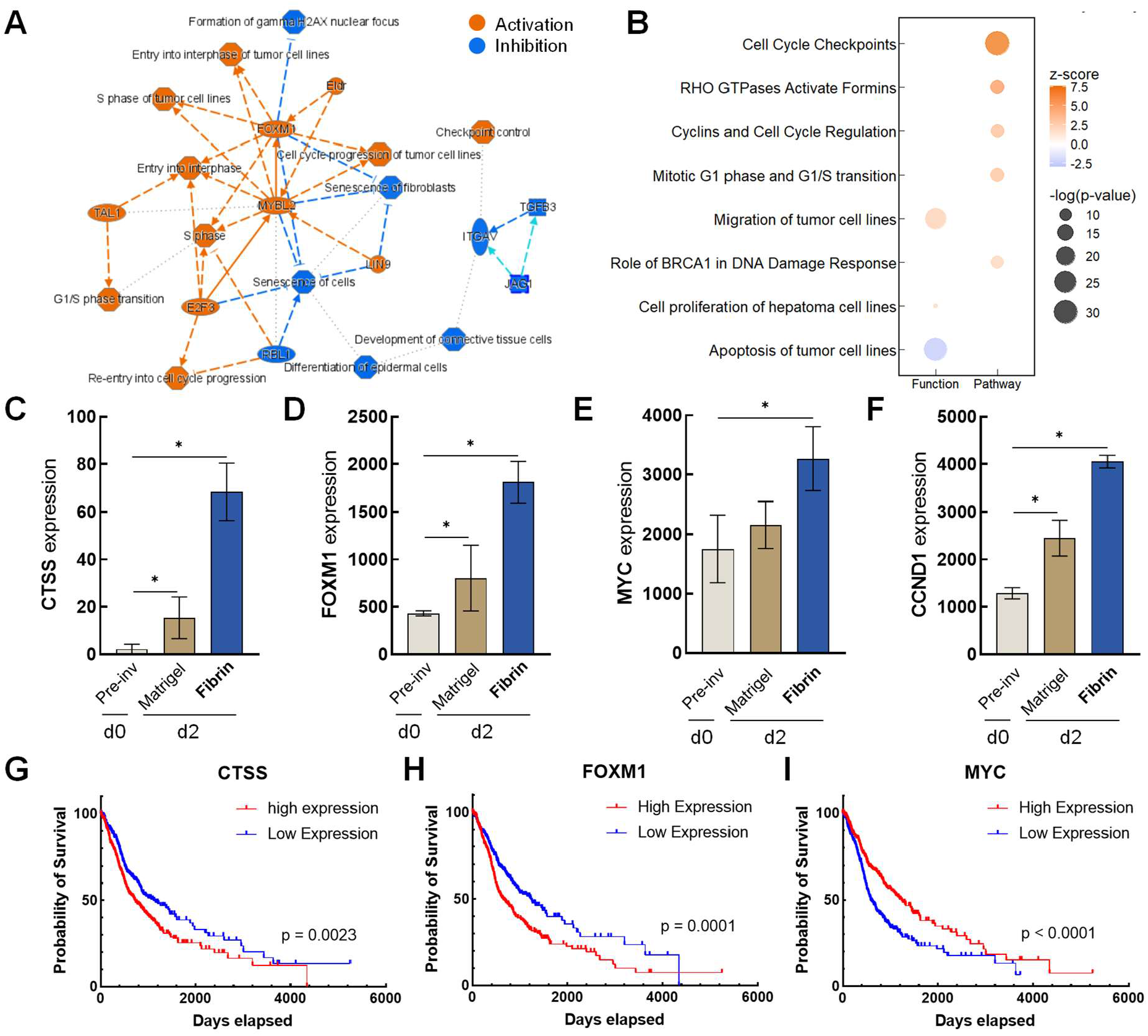
Functional enrichment and clinical survival-day analysis of GBM invasion-associated genes. **(A)** Network analysis of the DEGs in day 2 fibrin-embedded spheroids, and **(B)** top regulated pathways and upstream regulators identified by Ingenuity Pathway Analysis. (C–F) Normalized gene expression levels of **(C)** *CTSS*, **(D)** *FOXM1*, **(E)** *MYC*, and **(F)** *CCND1* across experimental conditions. (G–I) Kaplan-Meier survival analysis based on mRNA expression levels of **(G)** *CTSS*, **(H)** *FOXM1*, and **(I)** *MYC* in LGG and GBM patients from the TCGA database (n=669). Statistically significant differences are indicated (\**p* < 0.05 by the Mann-Whitney U test).

### 2.3. A 3D GBM invasion platform for identifying potential repurposable drug candidates

To validate the utility of our 3D GBM invasion platform for drug testing, we evaluated the effects of four licensed drugs (dinaciclib, ixazomib, triptolide, and napabucasin) that are not currently indicated for GBM, along with temozolomide (TMZ), the FDA-approved first-line chemotherapy for GBM. The four drugs were selected based on the following procedure and rationale: a) we ranked drugs based on anti-proliferation (viability) results from drug repurposing screening of the PRISM database by Corsello et al.^[22]^ which encompasses 4,518 drugs tested against 578 human cancer cell lines. We focused on effects of these drugs on 17 GBM cell lines and supplementary Figure 1 represents the most potent 14 licensed drugs, not indicated for GBM, that exhibit at least a four-fold or greater reduction in viability across all 17 GBM cell lines in 2D culture compared to untreated controls; all 14 drugs demonstrate more potent anti-proliferation effects than TMZ.^[22]^ b) Considering TMZ’s low molecular weight (MW) (194 g/mol) and its ability of crossing BBB (lipophilicity), we selected four drugs that have small MW (< 400 g/mol) and high lipophilicity (log of octanol-water partition coefficient: 1.3 < log*P* < 2.6). These physical properties likely enhance the ability of drugs to passively penetrate the BBB due to their impact on cellular permeation and membrane affinity.^[24]^

We assessed effects of the four drugs (dinaciclib, ixazomib, triptolide, and napabucasin) on growth, invasion, and morphology of GBM spheroids in the biomimetic 3D fibrin scaffold, and compared them to TMZ **(Figure 4)**. The GBM spheroids were treated for 48 hours before stained with the live-cell fluorescent viability dye Calcein-AM and imaged using a spinning disk confocal microscope. The images were binarized based on signal intensity. The morphological metrics, area and border complexity, were measured based on the binarized contour (Figure 4). Untreated control (0.1% DMSO) and TMZ (both 1 and 10 µM)-treated GBM spheroids developed many long protrusions, forming complex boundaries **(Figure 4A)**.

**Figure 4.**
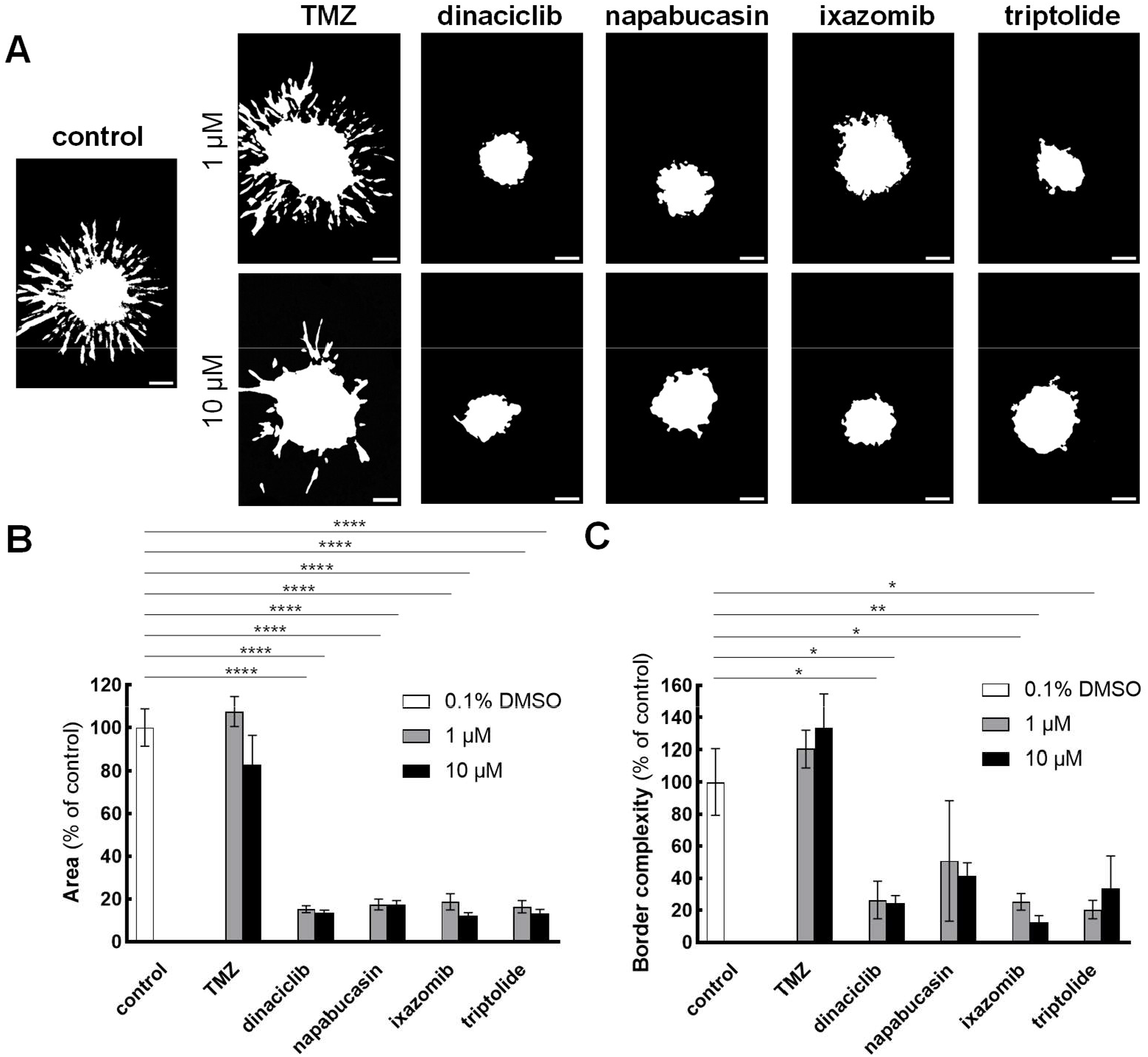
Evaluation of the anti-proliferative and anti-invasive effects of the selected four drugs as potential repurposable candidates in 3D GBM invasion platform. **(A)** Representative binarized live-cell images of GBM spheroids embedded in fibrin scaffolds and treated for 48 h with 0.1% DMSO (control), temozolomide (TMZ), or four selected repurposed drugs (dinaciclib, napabucasin, ixazomib, triptolide) at 1 µM and 10 µM concentrations. Scale bars: 200 µm. Quantitative analysis of **(B)** total spheroid area and **(C)** border complexity, expressed as a percentage of the control. Statistically significant differences relative to the control are indicated (\**p* < 0.05, \*\**p* < 0.01, \*\*\*\**p* < 0.0001) as determined by one-way ANOVA. Data represent mean ± SEM (n = 5 independent replicates).

We did not observe significant differences between control and TMZ treated groups in cell growth area **(Figure 4B)** and border complexity **(Figure 4C)**. In contrast, all four drugs (dinaciclib, napabucasin, ixazomib, and triptolide) at both 1 and 10 µM significantly inhibited growth area (**Figure 4B**). Dinaciclib and ixazomib at both 1 µM and 10 µM, as well as triptolide at 1 µM, significantly inhibited border complexity. Although triptolide at 10 µM and napabucasin at both concentrations did not reach statistical significance, it still demonstrated an inhibitory trend in border complexity (**Figure 4C**). Though actual images of the spheroid boundary morphologies exhibit drastic differences between untreated and all four drug-treated groups (**Figure 4A**), statistical differences in border complexity (\**p*<0.05 and **p < 0.01) are smaller (i.e. higher variations (SEM)) than those of growth area (\*\*\*\**p* <0.0001). This suggests that the complex and variable shapes of the boundaries of GBM spheroids are not accurately captured with high sensitivity using a traditional border complexity formula, defined as inverse of circularity 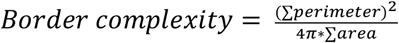^[25]^. To address the limitation of simplified formulas for quantifying morphological changes (*See Discussion for details*), we have applied deep-learning (DL) enabled high-content morphodynamic analysis to accurately quantify and comprehensively capture the morphodynamics of GBM spheroids in a 3D ECM mimetic platform, which are presented in **Figure 5** and **Supplementary Figures 2 to 4**.

**Figure 5.**
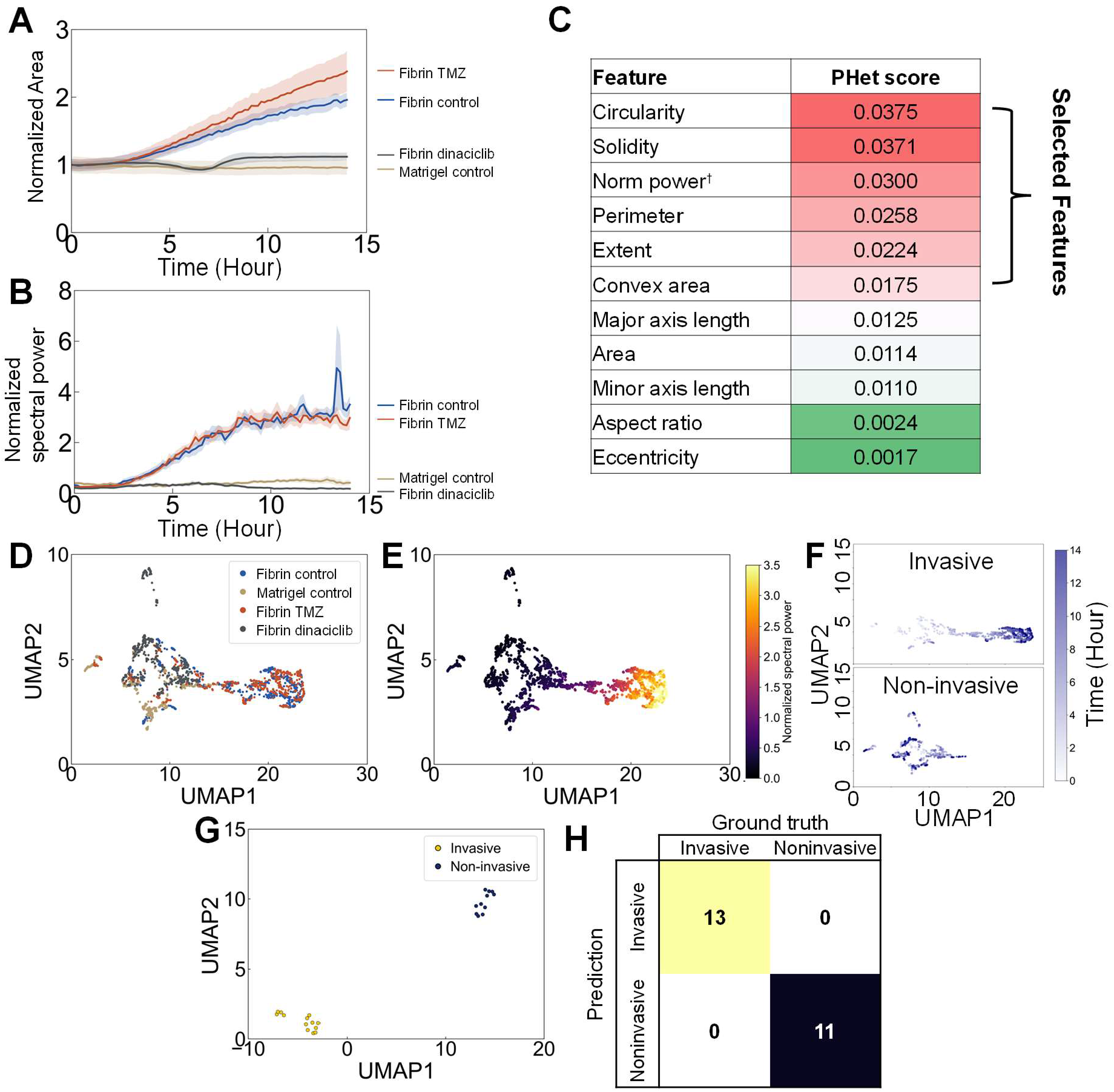
Deep-learning-enabled high-content morphodynamic analysis and predictive modeling of GBM invasion. Longitudinal quantification of **(A)** normalized spheroid area and **(B)** normalized spectral power for spheroids across different experimental conditions over 14 hours. **(C)** Ranking of morphodynamic features using the PHet algorithm to prioritize “Heterogeneity-preserving Discriminative” (HD) features that maximize group separation while maintaining biological diversity. **(D)** UMAP embedding of the top six prioritized features for all captured frames, demonstrating condition-specific distribution patterns. **(E)** UMAP visualization color-coded by normalized spectral power, showing a transition from low-complexity (purple) to high-complexity (yellow) phenotypes along the first manifold axis. **(F)** Representative temporal trajectories of invasive and non-invasive spheroids through the UMAP morphological state space. **(G)** UMAP clustering based on the top six prioritized features, with each feature individually averaged over the 7 frames (10 min intervals) covering the 8th hour of live-cell imaging. **(H)** Confusion matrix of prediction model demonstrating 100% predictive accuracy achieved by the classifier using the 8th-hour averaged features to forecast the terminal invasive phenotype.

### 2.4. A deep learning pipeline enables high-content morphodynamic analysis of GBM invasion

To overcome the limitations of traditional simple quantification metrics, we implemented a high-content analysis pipeline centered on a DL algorithm MARS-Net for robust and accurate segmentation of complex spheroid boundaries from live-cell imaging data **(Figure 5)**. This automated workflow enabled the extraction of a rich set of multi-parametric morphodynamic features that describe the area and shape of the invading spheroids **(Figure 1A-1B; Supplementary Figure 2)**. To quantitatively characterize morphological complexities beyond area and shape descriptors, we performed Fourier transformation of the spheroid boundary, extracting a total of 341 features describing shape, area, and the PCA dimension reduction and the spectral components of the Fourier transformed contour.

A key component of this pipeline is the calculation of spectral power, which serves as a sensitive indicator of boundary complexity of invading spheroids. The spectral power array represents the spectral components (*P*_*k*_) obtained through a Fourier transform of the spheroid’s boundary coordinates, providing a translation- and rotation-invariant phenotype of spheroid boundary morphology that is robust and scale invariant (independent of spheroid sizes). Furthermore, spectral approaches allow for targeted denoising of the higher-frequency components often associated with segmentation noise (pixelation/threshold jitter) based on frequency. This makes the metric more robust across different microscopes and segmentation pipelines and improves comparability across experiments. In contrast, traditional scalar morphometrics (e.g., inverse circularity, form factor) compress boundary geometry into a single value, and could map distinct boundary morphologies (e.g., a spheroid with a single long protrusion versus one with many tiny protrusions) to the same metric value, obscuring critical differences in invasive shape pattern. To provide a direct indicator of overall invasiveness and boundary complexity, we utilized normalized spectral power, defined as the scalar sum of the components within the spectral power array. A higher sum indicates a more convoluted, irregularly shaped boundary, while a lower sum corresponds to a smoother, weakly invasive phenotype **(Figure 5B)**.

### 2.5. High-dimensional feature analysis sensitively distinguishes different invasive phenotypes

We next applied a high-content analysis pipeline to distinguish the nuanced effects of therapeutic agents on GBM invasion dynamics. Initial dimensional reduction analysis of all live-cell images using the full feature set resulted in overlapping UMAP clusters **(Supplementary Figure 3)**, which indicates that non-discriminative variance dominated the global features.

To address this, we employed the Preserving Heterogeneity (PHet) algorithm to identify the most informative features that maintain within-class heterogeneity while maximizing inter-class discrimination **(Figure 5C)**. PHet prioritized the subtle morphological and structural irregularities specifically associated with aggressive invasive phenotypes.^[21]^ Refining the analysis to the top six PHet-scored features led to clear distinctions between highly invasive and non-invasive spheroid boundaries. UMAP dimensional reduction based on the six features revealed that the more invasive spheroids (untreated control and TMZ-treated spheroids in fibrin) and less-invasive spheroids (dinaciclib-treated spheroids in fibrin and spheroids in Matrigel) demonstrated distinctive distribution patterns **(Figure 5D)**. While an invasive spheroid transits from the lower normalized spectral power region to the higher spectral power region as the time increases, a non-invasive spheroid remains in the low spectral power region throughout the imaging period **(Figure 5 E and F)**. This finding demonstrates that the high-content morphodynamic profiling enables accurate, sensitive, and automated digitalization of GBM spheroid invasion.

### 2.6. Enabling early detection of invasive phenotypes

We investigated the prediction ability of early invasive phenotypes. While the rapid turnaround time is a primary advantage of 3D *in vitro* drug testing platforms, not all tumor organoid types exhibit substantial invasion within the first few days. Depending on the cancer types and cell source, tumor spheroid/organoid models may require days or weeks to demonstrate invasiveness. Therefore, predicting long-term tumor invasiveness from early-stage live-cell imaging offers a significant advantage by shortening the time required for model disease progression and evaluate therapeutic strategies.

To achieve this, we analyzed consecutive images within one-hour windows at various time points. The six PHet-selected features were averaged within each window and processed using dimensional reduction-based clustering. As early as the eighth hour of live-cell imaging, invasive and non-invasive spheroids were clearly resolved into two distinct UMAP clusters **(Figure 5G, H)**. This high predictive accuracy and precision (>90%) were maintained throughout the subsequent imaging period, demonstrating that high-content morphodynamic analysis enables automated, objective, and early detection of invasive phenotypes. When validated using an independent experimental dataset, our model predicted invasive phenotypes with 95.2% precision and 83.3% accuracy using only the eighth hour images, confirming that this method is robust across experimental batches **(Supplementary Figure 4)**.

## 3. Discussion

In this study, we have developed an integrated platform that combines the fabrication of a 3D GBM invasion platform in an ECM-mimetic 3D fibrin scaffold coupled with a novel AI-driven pipeline for high-content,longitudinal morphodynamic analysis. Our work establishes a new paradigm for quantifying complex cell behaviors within engineered tissues, demonstrating a significant advance in the predictive power of 3D tumor invasion models.

A cornerstone of our platform is the use of the fibrin hydrogel, a departure from commonly used scaffolds such as Matrigel or hyaluronic acid, to mimic the hemorrhagic, pro-thrombotic perivascular niche of GBM. The compromised vasculature then leads to extravasation of plasma components, including fibrinogen, into the interstitial space. The stable polymerized fibrin network is a hallmark of high-grade gliomas that differs from the fibrin-free normal brain microenvironment.^[15]^ Previous studies have utilized fibrin to fabricate functional human brain microvascular networks within microfluidic devices, confirming their suitability for complex neuro-oncological biofabrication.^[26,27]^ Our results from transcriptomic profiling as well as morphodynamic analyses of cancer spheroids in real-time justify building the 3D GBM invasion model with fibrin scaffold. Future work can further advance this model to a vascularized coculture system recapitulating the GBM microenvironment tightly regulated by the BBB.

RNA-seq results confirm that this biomimetic fibrin scaffold is not merely a passive scaffold but an active modulator of the tumor phenotype. We demonstrate that the fibrin scaffold induces a profound shift towards a highly invasive morphology (Figure 1), which was correlated with a distinct transcriptomic signature characterized by the upregulation of cell cycle progression and migration pathways (Figure 2 and 3). We found that this phenotype is likely driven by the upregulation of *CTSS, CCND1, MYC*, and *FOXM1*, all are oncogenic drivers with expression levels positively correlated with the grade of glioma.^[28–35]^ Studies demonstrated that proliferation can promote cancer cell migration and invasion,^[36]^ though the relationship between the two is complex; some studies suggest that inhibiting proliferation can also inhibit invasion, others show that these processes can be regulated independently.^[37]^ All four target genes we have identified are directly or indirectly related to cell cycle, proliferation, and cancer cell invasion, while *CTSS* and *CCND1* are more directly related to cancer cell invasion.^[34,38,39]^ *CTSS* encodes Cathepsin S, a lysosomal cysteine protease that is critical for tumor establishment and local dissemination through extracellular matrix (ECM) remodeling.^[34,35]^ *CCND1*, which encodes Cyclin D1, regulates cell cycle transition by forming complexes with cyclin-dependent kinases (CDK4/6).^[40]^ While often localized in nuclei, CCND1/Cdk4 activity promotes GBM dissemination through mechanisms independent of the retinoblastoma protein (RB1) when localized to the cytoplasm.^[38]^ The *MYC* gene contributes to malignancy by aggressively driving proliferation, blocking differentiation, increasing cell migration, and inducing angiogenesis.^[41]^ *FOXM1* regulates various functions related to GBM malignancy, including uncontrolled proliferation, maintenance of cancer stemness, activating DNA damage response, and regulation of invasion by mediating *MMP2*.^[29,29,42]^ High *CTSS, MYC*, and *FOXM1* expressions predict poor survival outlook in glioma patients from the TCGA cohort (Figure 2G-I). Both *CTSS* and *FOXM1* mediate TMZ resistance^[42,34]^, which could partially explain the lack of efficacy when treating GBM spheroids in fibrin with TMZ (Figure 4). Taken together, our enriched gene pathways and the significant target genes identified from transcriptomic profiling results indicates that our ECM-mimetic fibrin scaffold creates a pro-invasive microenvironment for GBM spheroids by upregulating genes associated with increased cell proliferation and migration, ultimately contributing to tumor growth and invasion.

Using our RNA-seq data, we also examined interactions among genes involved in integrin-fibronectin (FN) axis as previous studies have identified the integrin-FN axis as the main route of fibrin-promoted GBM invasion. In GBM, FN is assembled into fibrillar structures and promotes the collective invasion of tumor cells into the basement membrane *in vitro* and *in vivo*.^[43]^ FN binds to αvβ3 and αvβ5 integrins on the cell membrane,^[43,44]^ activating downstream target focal adhesion kinase (FAK), thus promoting tumor growth and infiltration.^[45]^ However, in our sequencing data, *FN1, ITGAV, ITGB3, and ITGB5* were not significantly upregulated in GBM cultured in fibrin scaffold, suggesting that the enhanced invasion of GBM39 was regulated by a mechanism independent of the FN-integrin axis.

Traditional quantification methods, though widely used for cancer spheroids/organoids, distill complex morphologies into singular metrics such as area or circularity and are insufficient to capture the nuances of invasion. Although the anti-invasive effects of the four drugs were qualitatively evident in the live-cell images (Figure 4A), traditional formulas revealed significant analytical bottlenecks inherent in scalar metrics. While growth area provided a simple, representative indicator of therapeutic efficacy (Figure 4B), the border complexity metric (inverse circularity) exhibited high within-group variation and elevated standard error of the mean (SEM), particularly in the control untreated groups (Figure 4C). This instability frequently undermines statistical significance even when the biological inhibition of invasion is visibly apparent. We attribute this high variance to the ‘fuzziness’ of standard segmentation workflows based on the binarization of fluorescence signals. In 3D invasion assays, signal noise, pixelation artifacts, and the presence of loose cells at the leading edge can artificially inflate perimeter measurements, to which the border complexity metric is mathematically hypersensitive. These findings underscore that simple geometric ratios are insufficient for capturing the non-linear, high-dimensional nature of tumor invasiveness. The lack of discriminative power in these scalar features arises because conventional feature selection focuses solely on mean differences; this can oversimplify the feature space and stifle the characterization of biological heterogeneity in patient-derived GBM. To address this challenge, we present a robust DL-enabled segmentation method and high-content morphodynamic descriptors, which can filter experimental noise while preserving the subtle phenotypic variations essential for identifying effective personalized therapies.

Our work highlights the development and implementation of a DL-powered, high-dimensional analysis pipeline. While other studies have employed DL for segmentation, our approach is distinguished by its subsequent extraction and analysis of a rich set of morphodynamic features to create a “fingerprint” of invasion. This high-content approach enables us to uncover subtle therapeutic effects that were completely invisible to standard metrics. Moreover, a key advantage of this integrated platform is its ability to leverage longitudinal data to enhance predictive accuracy. The automated and accurate classification achieved using dimensional reduction of PHet-selected feature (Figure 5) proves that the temporal evolution of a spheroid’s morphology is a far more comprehensive and robust identifier of its phenotype than the endpoint measurement based on a single frame of capturing. Building on this, we demonstrate that our model can predict a spheroid’s long-term invasive fate with high accuracy and precision using only partial image sets from early time points (e.g., the eighth hours), rather than requiring the full time-course images through the terminal end point (Figure 5 (14 hours); Supp Fig 4 (12 hours)). This is a critical advance for the field of *in vitro* drug testing. Many patient-derived organoid or complex tissue models exhibit slow-developing phenotypes that can take weeks to become apparent. Our finding suggests that these prolonged experiments could be dramatically accelerated, enabling higher-throughput testing and faster decision-making in drug development pipelines. This predictive capability transforms the platform from a descriptive tool into a prognostic one.

Unlike conventional 2D cell culture, our platform provides an ECM-mimetic 3D fibrin scaffold, a key feature of the tumor microenvironment, and can serve as an *in vivo*-like pre-clinical model for studying how ECM composition affects cancer invasion and for evaluating chemotherapeutic agents. Moreover, the innate suitability of fibrin in modeling angiogenesis opens the future potentials for incorporating our 3D GBM invasion model into advanced microfluidic GBM-on-a-chip systems with a BBB^[26,27]^; this will enable to investigate the interplay of immune and stromal cells present in GBM TME. Future studies can further classify morphodynamic signatures we have identified for GBM spheroid invasiveness and responses to anti-invasive drugs, by regulating cell adhesion (e.g., Cilengitide) and cytoskeletal pathways, and the target genes we have identified.

## 4. Conclusion

In summary, we have developed a 3D GBM invasion platform that recapitulates the hemorrhagic, pro-thrombotic extracellular matrix of glioblastoma using a fibrin scaffold. This engineered microenvironment induces a pro-invasive and pro-proliferative phenotype in PDX GBM spheroids, by upregulating key oncogenic cell cycle regulators and ECM-remodeling proteases. We have established a platform to digitally and accurately quantify tumor invasion and aggressiveness-related phenotypes and predict long-term invasive phenotypes from early-stage morphological changes, by integrating this *in vivo*-like invasion model with MARS-Net, a deep-learning (DL) segmentation pipeline, and high-content morphodynamic analysis. The DL quantitative approach overcomes the limitations of traditional scalar metrics, enabling the accurate evaluation of anti-invasive therapeutics. Our 3D platform offers a robust and comprehensive, quantitative approach that accelerates high-throughput drug screening and serves as a foundation for discovering novel anti-invasive agents.

## 5. Materials & methods

### 5.1. Culture of patient-derived xenograft (PDX) glioblastoma (GBM) cells

PDX-derived GBM cell lines #39 and #59 were originally obtained from the Sarkaria lab under material transfer agreement (Mayo Clinic, Rochester, MN).^[46]^ PDX tissues were maintained and serially passaged in nude mice before isolation for cell and spheroid culture.^[47]^ Primary PDX cells were cultured in Dulbecco’s Modified Eagle Medium (DMEM, Gibco, Waltham, MA) supplemented with 10% Fetal Bovine Serum (Gibco, Waltham, MA) and 1% Penicillin-Streptomycin (Gibco, Waltham, MA). All cells were maintained in a humidified incubator at 37°C and 5% CO_2_.

### 5.2. GBM spheroid formation

Single-cell suspensions were prepared from confluent cultures using TrypLE Express (Gibco, Waltham, MA). To form spheroids, cells were seeded at a density of 500,000 cells/well in 1.5 mL of culture medium into AggreWell 800 24-well plate (STEMCELL Technologies, Cambridge, MA). The plates were centrifuged at 100 g for 5 minutes to facilitate cell aggregation and incubated overnight to allow for the formation of compact, uniform spheroids with an approximate diameter of 150-200 µm.

### 5.3. Hydrogel preparation and spheroid embedding

Two distinct hydrogel matrices — fibrin and Matrigel— were prepared to investigate 3D GBM spheroid invasion.

#### Fibrin hydrogel

Fibrinogen (Sigma-Aldrich, St. Louis, MO) was dissolved into a final concentration of 50 mg/mL. Thrombin (Sigma-Aldrich, St. Louis, MO) was prepared at a concentration of 100 U/mL. Pre-formed spheroids were gently mixed with the fibrinogen solution, and 50 µL of the spheroid-fibrinogen suspension containing a final concentration of 3 mg/mL fibrinogen was dispensed into each well of a 96-well glass-bottom plate. Polymerization was initiated by adding 1 µL of the thrombin solution to each well and incubating at 37°C for 15 minutes.

#### Matrigel

Growth factor reduced basement membrane matrix (Matrigel, Corning, Corning, NY) was thawed at 4°C overnight. Pre-formed spheroids were suspended in cold culture medium and mixed with Matrigel at a 1:1 ratio for a final concentration of 5 mg/mL. A volume of 50 µL of the spheroid-Matrigel suspension was added to pre-chilled 96-well plates and allowed to polymerize at 37°C for 30 minutes. Following polymerization, 150 µL of complete culture medium was added atop each hydrogel.

### 5.4. Identification of the four candidate drugs from the PRISM drug repurposing screening database

The following procedures were applied for selecting the four drugs to be tested for GBM spheroids in the biomimetic 3D fibrin scaffold: First, drugs were ranked based on their anti-proliferative (anti-viability) effect results from drug repurposing screening of the PRISM database (https://depmap.org/repurposing); specifically dataset by Corsello et al.^[22]^, which encompasses 4,518 drugs tested against 578 human cancer cell lines, was used. Drugs demonstrating at least a four-fold or greater reduction in viability across all 17 GBM cell lines in conventional 2D culture compared to untreated controls were selected. The 14 candidate drugs were ranked based on their lowest median log_10_ percent viability compared to control groups across the range of tested doses, while TMZ was included as a reference for comparison with these drugs’ effects (Supplementary Figure 1). Second, for each of these compounds, key molecular and clinical parameters were documented, including MW, mechanism of action, and known primary disease indication. Logarithm of octanol-water partition coefficient (log*P*) values were obtained as experimental data from DrugBank^[48]^ or as predicted values via ChemAxon (http://www.chemaxon.com), where experimental data was unavailable. Then, among the 14 drugs, four were chosen based on their small MW (< 400 g/mol), and high lipophilicity (1.3 < log*P* < 2.6): dinaciclib (396 g/mol), napabucasin (240 g/mol), ixazomib (361 g/mol), and triptolide (360 g/mol).

### 5.5. 3D spheroid invasion assay and drug treatment

The drugs tested in this study were: temozolomide (TMZ), dinaciclib, ixazomib, triptolide, and napabucasin (MedChemExpress, Princeton, NJ).These drugs were dissolved in dimethyl sulfoxide (DMSO) (Sigma-Aldrich, St. Louis, MO) at stock concentrations 100 mM (TMZ, dinaciclib, and ixazomib), 80 mM (triptolide), and 20mM (napabucasin), based on their maximal solubility, and stored at -80ºC in the dark. Final drug concentrations (1 µM and 10 µM) were prepared by diluting the stock solutions in culture medium, ensuring the final DMSO concentration at ≤ 0.1%. Drug-containing medium was added to the wells immediately after spheroid embedding. Control wells received medium containing 0.1% DMSO. At the end of the treatment, the spheroids were stained with Calcein AM live-cell stain (Invitrogen, Waltham, MA) and imaged using a spinning disk confocal microscope (Yokogawa Electric Corporation of America, Houston, TX). To track spheroid contours, images were binarized based on the fluorescence emission signal (approximately 517 nm).

### 5.6. Time-lapse microscopy

Spheroid invasion was monitored using a Nikon Eclipse TiE inverted microscope (Nikon, Melville, NY) equipped with a 10x objective and an environmental chamber maintained at 37°C and 5% CO_2_. Spheroids were imaged via Differential Interference Contrast (DIC) every 10 minutes for 12 hours (Supplementary Figures 4) or 14 hours (Figure 5).

### 5.7. Bulk RNA sequencing and analysis

GBM39 spheroids collected immediately after formation served as the pre-invasive control, whereas the spheroids cultured in either fibrin or Matrigel were collected according to the experimental period (48 hours). Total RNA was extracted using the AllPrep DNA/RNA/Protein Mini Kit (QIAGEN, Louisville, KY) according to the manufacturer’s instructions. Bulk mRNA sequencing (RNA-seq) library preparation and sequencing were performed by Psomagen (Rockville, MD). Library preparation was performed using the TruSeq stranded mRNA kit and sequencing was conducted on NovaSeqX to generate 30 million paired-end reads per sample.

Raw RNA-seq reads were preprocessed with TrimGalore (https://www.bioinformatics.babraham.ac.uk/projects/trim_galore/) to remove reads with mean sequence quality below quality score 30, and to clip 13 bases from 5’ and 2 bases from 3’ ends of the reads. The reads were then aligned to the Ensembl Transcriptomes v96 using Kallisto.^[49]^ Differential gene expression analysis was performed using DESeq2 in R, with a significance threshold of an adjusted p-value < 0.05.^[50]^ Pathway and network analyses were conducted using Ingenuity Pathway Analysis (QIAGEN, Redwood City, CA).

### 5.8. TCGA survival-day analysis

The mRNA expression levels of low-grade glioma (LGG) and GBM patients were downloaded from the Cancer Genome Atlas (TCGA) database (https://www.tcga-survival.com). For each of our identified genes (CTSS, FOXM1, MYC, and CCND1), their expression levels were stratified into low and high expression groups based on the median expression threshold using GraphPad Prism version 10 (GraphPad Software, San Diego, CA). Kaplan Meier survival analysis was then conducted to determine the correlation between gene expression and patient survival-days.

### 5.9. High-content image analysis pipeline

A custom computational pipeline was developed to quantify spheroid morphodynamics from the time-lapse image series.

#### Spheroid segmentation

Spheroid boundaries were segmented from brightfield images using MARS-Net, a deep-learning-based pipeline, as we described.^[20]^ A manual segmentation (10 percent of the entire dataset) was performed to generate ground-truth annotations. These annotations were used to train the MARS-Net model until a loss less than 0.05 and a dice coefficient greater than 0.95 was achieved.

#### Morphodynamic feature extraction

From the segmented masks, a high-dimensional set of 341 morphodynamic features was extracted for each spheroid at each timepoint. This feature set consists of 13 region property features, normalized spectral power (see ‘*Spectral power calculation’* below), 200 features based on principle component analysis (PCA) dimension reduction of the Fourier transformed contour, and 127 features based on the spectral components of the Fourier transformed contour. The 13 region property features were categorized into descriptors of size (area, perimeter, major axis length, minor axis length, convex area), shape (circularity, solidity, eccentricity, extent, aspect ratio, orientation), and texture/complexity (mean intensity, intensity standard deviation).

#### Spectral power calculation

The spectral power is a spectrum-based measurement of boundary complexity that retains the full contour information prior to summarization. This was used so that distinct invasive morphologies would be distinguishable even when traditional single-value metrics (e.g., inverse circularity) would overlap. Specifically, individual organoid boundaries, which were segmented in 2D images to yield an ordered set of boundary individual organoid boundaries were segmented in 2D images to yield an ordered set of boundary coordinates 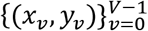 with periodic indexing (*x*_−1_, *y*_−1_ ) ≡ (*x*_*V*−1_, *y* _*V*−1_). The total contour length was computed as:

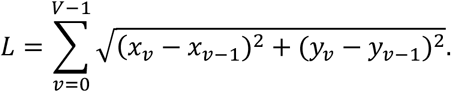

To enable Fourier analysis with uniform sampling, the boundary was re-parameterized by arc length and resampled to *M* equally spaced points 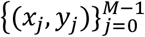 at spacing *L*/*M* along the contour (here *M* was fixed across samples, e.g., *M* = 256). The discrete Fourier transform was applied separately to the *x* and *y* coordinate sequences. For *x*, the transformation was defined as:

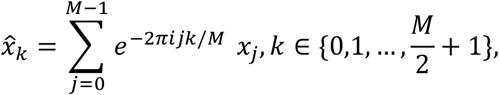

with an analogous definition for ŷ_k_. Spectral power at frequency *k*was computed as:

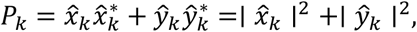

where “*” denotes complex conjugation. The *k* = 0 term (translation/centroid component) was excluded to enforce translation invariance, and use of squared magnitudes provides invariance to in-plane rotation. To obtain a single scalar phenotype that is scale-invariant and emphasizes higher-frequency boundary features, the spectrum was normalized by the first mode *P* and a curvature-weighting transform was applied. The resulting weighted spectral power was defined as:

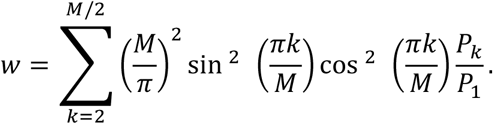

Weighted spectral power was computed only for contours with *M* ≥ 4 (even *M*) and non-zero normalization *P* > 0, and after enforcing periodic closure of the resampled boundary.^[51]^

#### Feature prioritization and dimensionality reduction

To identify the most important features for distinguishing invasive phenotypes, the Preserving Heterogeneity (PHet) algorithm was applied.^[21]^ The top-ranked features were then used as input for Uniform Manifold Approximation and Projection (UMAP) to visualize the high-dimensional data in a 2D space.

### 5.10. Statistical analysis

Data are presented as mean ± standard deviation (SD) or standard error of the mean (SEM) from at least three independent experiments, unless otherwise noted. Statistical significance between two groups was determined using a non-parametric Mann-Whitney U test, as indicated in each figure legend. For comparisons involving more than two groups, a one-way ANOVA was performed, followed by post-hoc multiple comparisons tests. A *p-value* < 0.05 was considered statistically significant. All statistical analyses were performed using GraphPad Prism version 10.1.0. (GraphPad Software, San Diego, CA).

## Acknowledgments

This work was supported by grants from the National Cancer Institute (NCI) R21CA220111 (to EHA) and R01CA279948 (to EHA) of the National Institutes of Health (NIH); The Sol Goldman Pancreatic Cancer Research Center Grant (to DHK) of the Johns Hopkins University. The content is solely the responsibility of the authors and does not necessarily represent the official views of the NIH. Authors thank Asmitha Sathya for TCGA gene expression and survival-day analysis; Professors John Laterra and Betty Tyler for accessing GBM39 and GBM59 cell lines.

## Data Availability Statement

The data that support the findings of this study are available from the corresponding author upon reasonable request.

## References

1. Louis DN, Perry A, Reifenberger G, von Deimling A, Figarella-Branger D, Cavenee WK, et al. The 2016 World Health Organization Classification of Tumors of the Central Nervous System: a summary. Acta Neuropathol 2016;131:803–20. doi: 10.1007/s00401-016-1545-1.

2. Paw I, Carpenter RC, Watabe K, Debinski W, Lo H-W. Mechanisms regulating glioma invasion. Cancer Letters 2015;362:1–7. doi: 10.1016/j.canlet.2015.03.015.

3. Mair DB, Ames HM, Li R. Mechanisms of invasion and motility of high-grade gliomas in the brain. Mol Biol Cell 2018;29:2509–15. doi: 10.1091/mbc.E18-02-0123.

4. Lee SY. Temozolomide resistance in glioblastoma multiforme. Genes Dis 2016;3:198–210. doi: 10.1016/j.gendis.2016.04.007.

5. Bhowmik A, Khan R, Ghosh MK. Blood brain barrier: a challenge for effectual therapy of brain tumors. Biomed Res Int 2015;2015:320941. doi: 10.1155/2015/320941.

6. Pu J, Yuan K, Tao J, Qin Y, Li Y, Fu J, et al. Glioblastoma multiforme: an updated overview of temozolomide resistance mechanisms and strategies to overcome resistance. Discov Onc 2025;16:731. doi: 10.1007/s12672-025-02567-3.

7. Liu P, Griffiths S, Veljanoski D, Vaughn-Beaucaire P, Speirs V, Brüning-Richardson A. Preclinical models of glioblastoma: limitations of current models and the promise of new developments. Expert Rev Mol Med 2021;23. doi: 10.1017/erm.2021.20.

8. Sivakumar H, Devarasetty M, Kram DE, Strowd RE, Skardal A. Multi-Cell Type Glioblastoma Tumor Spheroids for Evaluating Sub-Population-Specific Drug Response. Front Bioeng Biotechnol 2020;8:538663. doi: 10.3389/fbioe.2020.538663.

9. Manduca N, Maccafeo E, De Maria R, Sistigu A, Musella M. 3D cancer models: One step closer to in vitro human studies. Front Immunol 2023;14. doi: 10.3389/fimmu.2023.1175503.

10. Xiao W, Sohrabi A, Seidlits SK. Integrating the glioblastoma microenvironment into engineered experimental models. Future Sci OA 2017;3:FSO189. doi: 10.4155/fsoa-2016-0094.

11. Xiao W, Wang S, Zhang R, Sohrabi A, Yu Q, Liu S, et al. Bioengineered scaffolds for 3D culture demonstrate extracellular matrix-mediated mechanisms of chemotherapy resistance in glioblastoma. Matrix Biol 2020;85–86:128–46. doi: 10.1016/j.matbio.2019.04.003.

12. Knowles LM, Kiwitt TC, Pilch J. Blood Clotting Contributes to a Malignant Glioma Phenotype: A Commentary. J Cancer Immunol 2025;7:123–7. doi: 10.33696/cancerimmunol.7.111.

13. Ruoslahti E. Brain extracellular matrix. Glycobiology 1996;6:489–92. doi: 10.1093/glycob/6.5.489.

14. Dzikowski L, Mirzaei R, Sarkar S, Kumar M, Bose P, Bellail A, et al. Fibrinogen in the glioblastoma microenvironment contributes to the invasiveness of brain tumor-initiating cells. Brain Pathol 2021;31:e12947. doi: 10.1111/bpa.12947.

15. Knowles LM, Wolter C, Ketter R, Urbschat S, Linsler S, Müller S, et al. Fibrin As a Target for Glioblastoma Detection and Treatment. Blood 2019;134:3630–3630. doi: 10.1182/blood-2019-124089.

16. Tehrani M, Friedman TM, Olson JJ, Brat DJ. Intravascular thrombosis in central nervous system malignancies: a potential role in astrocytoma progression to glioblastoma. Brain Pathol 2008;18:164–71. doi: 10.1111/j.1750-3639.2007.00108.x.

17. Sirenko O, Mitlo T, Hesley J, Luke S, Owens W, Cromwell EF. High-content assays for characterizing the viability and morphology of 3D cancer spheroid cultures. Assay Drug Dev Technol 2015;13:402–14. doi: 10.1089/adt.2015.655.

18. Vinci M, Box C, Zimmermann M, Eccles SA. Tumor spheroid-based migration assays for evaluation of therapeutic agents. Methods Mol Biol 2013;986:253–66. doi: 10.1007/978-1-62703-311-4_16.

19. Berens EB, Holy JM, Riegel AT, Wellstein A. A Cancer Cell Spheroid Assay to Assess Invasion in a 3D Setting. J Vis Exp 2015:53409. doi: 10.3791/53409.

20. Jang J, Wang C, Zhang X, Choi HJ, Pan X, Lin B, et al. A deep learning-based segmentation pipeline for profiling cellular morphodynamics using multiple types of live cell microscopy. Cell Rep Methods 2021;1:100105. doi: 10.1016/j.crmeth.2021.100105.

21. M. A. Basher Ar, Hallinan C, Lee K. Heterogeneity-preserving discriminative feature selection for disease-specific subtype discovery. Nat Commun 2025;16:3593. doi: 10.1038/s41467-025-58718-1.

22. Corsello SM, Nagari RT, Spangler RD, Rossen J, Kocak M, Bryan JG, et al. Discovering the anti-cancer potential of non-oncology drugs by systematic viability profiling. Nat Cancer 2020;1:235–48. doi: 10.1038/s43018-019-0018-6.

23. Pinzi L, Bisi N, Rastelli G. How drug repurposing can advance drug discovery: challenges and opportunities. Front Drug Discov 2024;4:1460100. doi: 10.3389/fddsv.2024.1460100.

24. Pajouhesh H, Lenz GR. Medicinal chemical properties of successful central nervous system drugs. NeuroRx 2005;2:541–53. doi: 10.1602/neurorx.2.4.541.

25. Hou Y, Konen J, Brat DJ, Marcus AI, Cooper LAD. TASI: A software tool for spatial-temporal quantification of tumor spheroid dynamics. Sci Rep 2018;8:7248. doi: 10.1038/s41598-018-25337-4.

26. Straehla JP, Hajal C, Safford HC, Offeddu GS, Boehnke N, Dacoba TG, et al. A predictive microfluidic model of human glioblastoma to assess trafficking of blood-brain barrier-penetrant nanoparticles. Proc Natl Acad Sci U S A 2022;119:e2118697119. doi: 10.1073/pnas.2118697119.

27. Lam MS, Aw JJ, Tan D, Vijayakumar R, Lim HYG, Yada S, et al. Unveiling the Influence of Tumor Microenvironment and Spatial Heterogeneity on Temozolomide Resistance in Glioblastoma Using an Advanced Human In Vitro Model of the Blood-Brain Barrier and Glioblastoma. Small 2023;19:e2302280. doi: 10.1002/smll.202302280.

28. Herms JW, von Loewenich FD, Behnke J, Markakis E, Kretzschmar HA. c-myc oncogene family expression in glioblastoma and survival. Surg Neurol 1999;51:536–42. doi: 10.1016/s0090-3019(98)00028-7.

29. Li Y, Zhang S, Huang S. FoxM1: a potential drug target for glioma. Future Oncol 2012;8:223–6. doi: 10.2217/fon.12.1.

30. Meng D, Chen Y, Yun D, Zhao Y, Wang J, Xu T, et al. High expression of N-myc (and STAT) interactor predicts poor prognosis and promotes tumor growth in human glioblastoma. Oncotarget 2015;6:4901–19. doi: 10.18632/oncotarget.3208.

31. Zhang D, Dai D, Zhou M, Li Z, Wang C, Lu Y, et al. Inhibition of Cyclin D1 Expression in Human Glioblastoma Cells is Associated with Increased Temozolomide Chemosensitivity. Cell Physiol Biochem 2018;51:2496–508. doi: 10.1159/000495920.

32. Zhang C, Yang M, Li Y, Tang S, Sun X. FOXA1 is upregulated in glioma and promotes proliferation as well as cell cycle through regulation of cyclin D1 expression. CMAR 2018; Volume 10:3283–93. doi: 10.2147/CMAR.S168217.

33. Mahzouni P, Taheri F. An Immunohistochemical Study of Cyclin D1 Expression in Astrocytic Tumors and its Correlation with Tumor Grade. Iran J Pathol 2019;14:252–7. doi: 10.30699/ijp.2019.82024.1771.

34. Fei M, Zhang L, Wang H, Zhu Y, Niu W, Tang T, et al. Inhibition of Cathepsin S Induces Mitochondrial Apoptosis in Glioblastoma Cell Lines Through Mitochondrial Stress and Autophagosome Accumulation. Front Oncol 2020;10:516746. doi: 10.3389/fonc.2020.516746.

35. Wei L, Shao N, Peng Y, Zhou P. Inhibition of Cathepsin S Restores TGF-β-induced Epithelial-to-mesenchymal Transition and Tight Junction Turnover in Glioblastoma Cells. J Cancer 2021;12:1592–603. doi: 10.7150/jca.50631.

36. Massaro RR, Faião-Flores F, Rebecca VW, Sandri S, Alves-Fernandes DK, Pennacchi PC, et al. Inhibition of proliferation and invasion in 2D and 3D models by 2-methoxyestradiol in human melanoma cells. Pharmacological Research 2017;119:242–50. doi: 10.1016/j.phrs.2017.02.013.

37. O’Connell MP, Marchbank K, Webster MR, Valiga AA, Kaur A, Vultur A, et al. Hypoxia Induces Phenotypic Plasticity and Therapy Resistance in Melanoma via the Tyrosine Kinase Receptors ROR1 and ROR2. Cancer Discovery 2013;3:1378–93. doi: 10.1158/2159-8290.CD-13-0005.

38. Cemeli T, Guasch-Vallés M, Nàger M, Felip I, Cambray S, Santacana M, et al. Cytoplasmic cyclin D1 regulates glioblastoma dissemination. The Journal of Pathology 2019;248:501–13. doi: 10.1002/path.5277.

39. Body S, Esteve-Arenys A, Miloudi H, Recasens-Zorzo C, Tchakarska G, Moros A, et al. Cytoplasmic cyclin D1 controls the migration and invasiveness of mantle lymphoma cells. Sci Rep 2017;7:13946. doi: 10.1038/s41598-017-14222-1.

40. Sun T, Xu Y-J, Jiang S-Y, Xu Z, Cao B-Y, Sethi G, et al. Suppression of the USP10/CCND1 axis induces glioblastoma cell apoptosis. Acta Pharmacol Sin 2021;42:1338–46. doi: 10.1038/s41401-020-00551-x.

41. Tateishi K, Iafrate AJ, Ho Q, Curry WT, Batchelor TT, Flaherty KT, et al. Myc-Driven Glycolysis Is a Therapeutic Target in Glioblastoma. Clin Cancer Res 2016;22:4452–65. doi: 10.1158/1078-0432.CCR-15-2274.

42. Zhang N, Wu X, Yang L, Xiao F, Zhang H, Zhou A, et al. FoxM1 inhibition sensitizes resistant glioblastoma cells to temozolomide by downregulating the expression of DNA-repair gene Rad51. Clin Cancer Res 2012;18:5961–71. doi: 10.1158/1078-0432.CCR-12-0039.

43. Serres E, Debarbieux F, Stanchi F, Maggiorella L, Grall D, Turchi L, et al. Fibronectin expression in glioblastomas promotes cell cohesion, collective invasion of basement membrane in vitro and orthotopic tumor growth in mice. Oncogene 2014;33:3451–62. doi: 10.1038/onc.2013.305.

44. Knowles LM, Gurski LA, Maranchie JK, Pilch J. Fibronectin Matrix Formation is a Prerequisite for Colonization of Kidney Tumor Cells in Fibrin. J Cancer 2015;6:98–104. doi: 10.7150/jca.10496.

45. Knowles LM, Wolter C, Linsler S, Müller S, Urbschat S, Ketter R, et al. Clotting Promotes Glioma Growth and Infiltration Through Activation of Focal Adhesion Kinase. Cancer Research Communications 2024;4:3124–36. doi: 10.1158/2767-9764.CRC-24-0164.

46. Vaubel RA, Tian S, Remonde D, Schroeder MA, Mladek AC, Kitange GJ, et al. Genomic and Phenotypic Characterization of a Broad Panel of Patient-Derived Xenografts Reflects the Diversity of Glioblastoma. Clin Cancer Res 2020;26:1094–104. doi: 10.1158/1078-0432.CCR-19-0909.

47. Rath P, Lal B, Ajala O, Li Y, Xia S, Kim J, et al. In Vivo c-Met Pathway Inhibition Depletes Human Glioma Xenografts of Tumor-Propagating Stem-Like Cells. Transl Oncol 2013;6:104–11. doi: 10.1593/tlo.13127.

48. Knox C, Wilson M, Klinger CM, Franklin M, Oler E, Wilson A, et al. DrugBank 6.0: the DrugBank Knowledgebase for 2024. Nucleic Acids Research 2024;52:D1265–75. doi: 10.1093/nar/gkad976.

49. Bray NL, Pimentel H, Melsted P, Pachter L. Near-optimal probabilistic RNA-seq quantification. Nat Biotechnol 2016;34:525–7. doi: 10.1038/nbt.3519.

50. Love MI, Huber W, Anders S. Moderated estimation of fold change and dispersion for RNA-seq data with DESeq2. Genome Biol 2014;15:550. doi: 10.1186/s13059-014-0550-8.

51. Padmanaban V, Tsehay Y, Cheung KJ, Ewald AJ, Bader JS. Between-tumor and within-tumor heterogeneity in invasive potential. Anderson ARA, ed. PLoS Comput Biol 2020;16:e1007464. doi: 10.1371/journal.pcbi.1007464.

