## Supplementary figures for "Deep-learning-enabled morphodynamic analysis of drug responses in a biomimetic fibrin-based 3D glioblastoma invasion model"

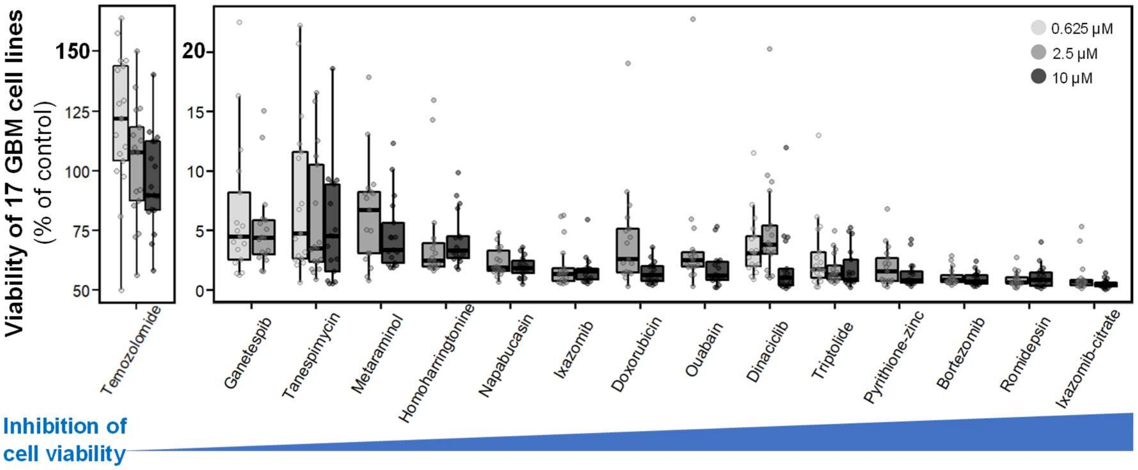


**Supplementary Figure 1.** Anti-proliferative effects of drugs were examined in GBM spheroids. Analysis on the PRISM drug repurposing resource (https://depmap.org/repurposing), which contains the anti-proliferation profile of 4,518 drugs tested on 578 human cancer cell lines, identified 14 drugs that inhibit the viability of 17 GBM cell lines by at least four-fold relative to control groups in 2D culture. The 14 drugs and TMZ at different doses were ranked by their lowest median $\log_{10}$ percent viability compared to the control. Original data is from the work by Corsello et al., 2020.

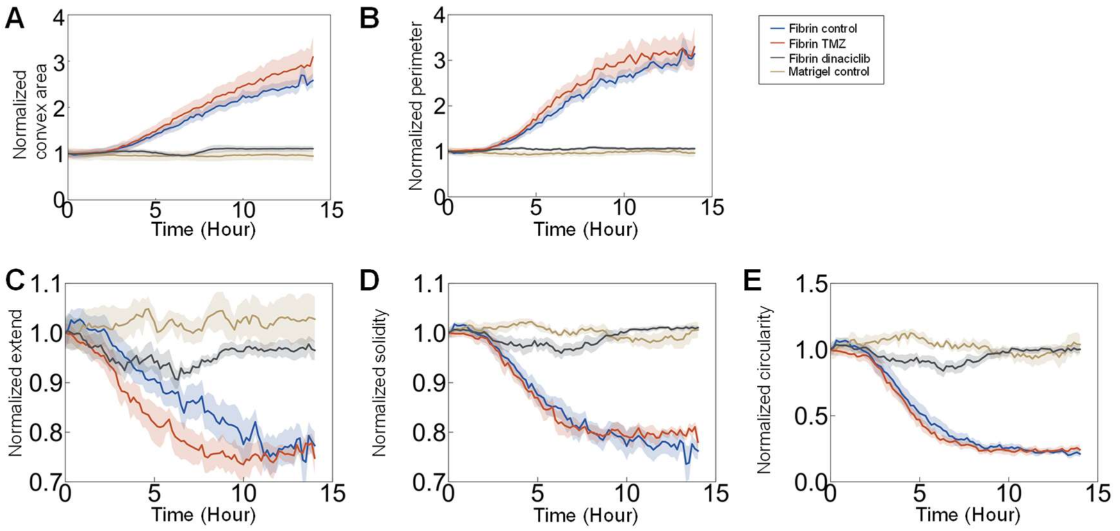


**Supplementary Figure 2**. **Real-time morphodynamic feature extraction using MARS-Net.** Longitudinal tracking of normalized morphological parameters over 14 hours. **(A)** Normalized convex area and **(B)** normalized perimeter exhibit a positive correlation with GBM spheroid invasion. Conversely, **(C)** normalized extent, **(D)** normalized solidity, and **(E)** normalized circularity are inversely correlated with the invasive phenotype. Data are presented in arbitrary units (A.U.) relative to the pre-invasive state.

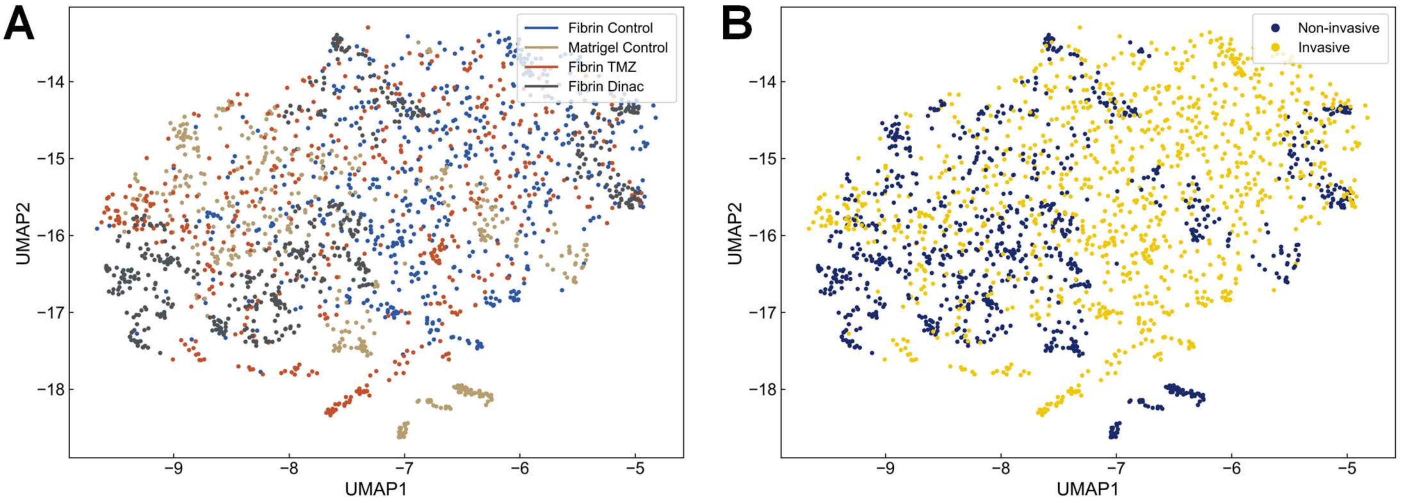


**Supplementary Figure 3**. **High-dimensional feature embedding of the global morphological landscape.** UMAP analysis utilizing the full feature set (N=341) annotated by **(A)** experimental condition and **(B)** invasive phenotype.

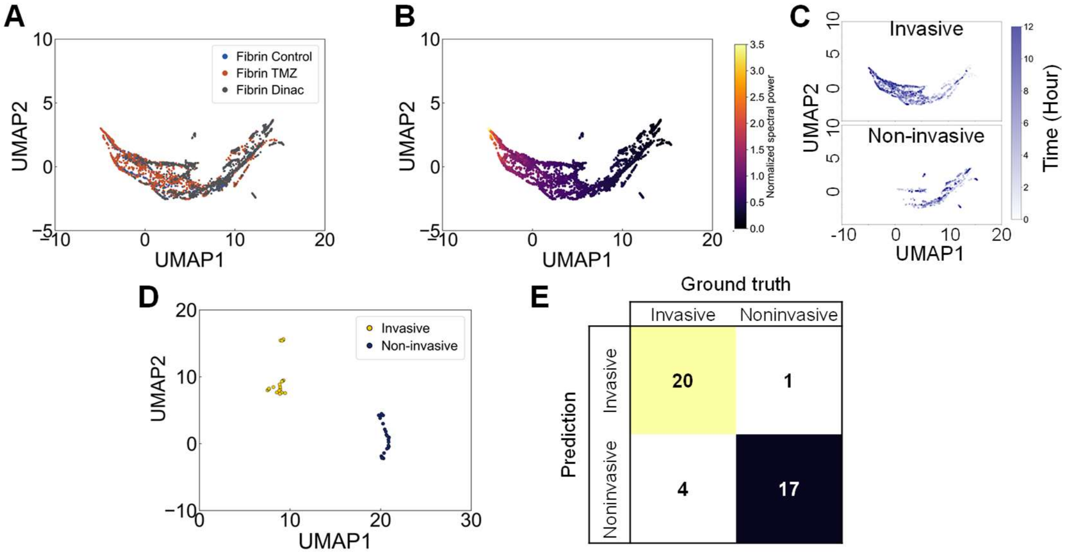


**Supplementary Figure 4. Independent validation of predictive morphodynamic analysis pipeline.** Analytical results of an independent live-cell imaging experiment demonstrate the reproducibility of the forecasting framework. UMAP embedding of the top six prioritized features for all captured frames annotated by **(A)** experimental condition and **(B)** normalized spectral power over 12 hours. **(C)** Representative temporal trajectories of invasive and non-invasive spheroids through the UMAP morphological state space. **(D)** UMAP clustering based on the top six PHet-prioritized features, with each feature value individually averaged over the 7 frames (10 min intervals) comprising the 8$^{th}$ hour of imaging. (E) Confusion matrix demonstrates 83.3% predictive accuracy and 95.2% precision in forecasting terminal invasive phenotypes using the 8$^{th}$-hour time-averaged signatures.
